# Capturing and Characterising Wild Yeast for Beer Brewing

**DOI:** 10.1101/2024.02.22.581081

**Authors:** Edward D. Kerr, Mark T. Howes, Benjamin L. Schulz

## Abstract

Beer is typically made using fermentation with *Saccharomyces cerevisiae* or *Saccharomyces pastorianus*, domesticated brewing yeasts. Historically, wild, non*-Saccharomyces* yeasts have also been frequently used in mixed culture fermentations to provide interesting and unique flavours to beer. However, brewing using mixed cultures or by spontaneous fermentation makes reproducing flavours and beer styles extremely difficult. Here, we describe a pipeline from collection of wild yeast from plant material to the characterisation and industrial scale production of beer using wild yeast. We isolated and identified wild yeast strains from the St Lucia campus of The University of Queensland, Brisbane, Australia. Several isolates fermented efficiently at laboratory scale, but failed to grow at industrial scale due to the combination of maltose and pressure stress. Systems biology showed that the synergistic metabolic defects caused by these dual stresses converged on amino acid nutrient uptake. Glucoamylase addition relieved maltose stress and allowed industrial scale fermentation using wild yeast. Our workflow allows efficient collection and characterisation of diverse wild yeast isolates, identification of interventions to allow their use at industrial scale, and investigation of the genetic and metabolic diversity of wild yeasts.

## Introduction

Fermentation is the key step in the beer brewing process. Fermentation with brewer’s yeast *Saccharomyces cerevisiae* is typically used to produce ales, while *S. pastorianus* is used for lager production. The inadvertent domestication of *S. cerevisiae* and hybridisation of *S. pastorianus* from *S. cerevisiae* and *S. eubayanus* by brewers eliminated the need to use spontaneous wild yeast inoculation for fermentation, and provided efficiency and predictability to brewing ^1,2^. Although efficiency and constancy ruled through this golden age of brewing, beers traditionally fermented with wild microbes were still made ^3,4^. It is well understood that the species and strain of yeast used in fermentation, with or without additional microbes, has a substantial impact on the flavour and sensory characteristics of the final beer ^5^. Spontaneous fermentation styles like lambics and American coolship ales (ACA), where wort is cooled and exposed to environmental microbes, can provide unique and unusual flavours and characteristics that are not seen in other styles of beer ^3,4^. A high species diversity of both yeast and bacteria are found throughout these spontaneous and mixed ferments ^3,4,6–8^. In contrast, few single strains of non-conventional yeasts have been investigated for beer production ^9^.

*Brettanomyces* yeast are the most common non-*Saccharomyces* yeast in the brewing and wine industry ^10–14^. They are often seen as spoilage yeasts in food and beverages, especially wine production ^10^. This association with spoilage is generally due to the flavours they produce: described as barnyard, medicinal, horse sweat, mousy, and metallic ^11–14^. In contrast, *Brettanomyces* are among the most commonly used non-*Saccharomyces* species in beer brewing, possessing desired traits that other non-*Saccharomyces* species lack ^15–17^. Many strains and species of *Brettanomyces* can utilise maltose, are ethanol tolerant, and can produce high amounts of ethyl caproate and ethyl caprylate, responsible for sweet, fruity, and pineapple or waxy, musty, and sweet flavours, respectively ^15–18^. In addition, several *Brettanomyces* isolates are able to hydrolyse maltotriose and other malto-oligosaccharides via extracellular alpha α-glucosidases ^19,20^.

*Torulaspora delbrueckii*, commonly described as a wine contaminant, is one of the most studied non-*Saccharomyces* yeasts ^21,22^ and its fermentation has been well-studied, particularly in wine production ^23–29^. Michel et al. (2016) screened *T. delbrueckii* strains collected from various sources including wine, cheese brine, wheat beer, and sorghum brandy using criteria including hop tolerance, ethanol production, amino acid and sugar utilisation, and sensory screening. All strains were found to be hop tolerant and require only limited amounts of amino acids ^22^. While most strains could use glucose, sucrose and, L-sorbose, only a few strains could efficiently use maltose ^21,22,30^. While many strains were only able to produce low quantities of ethanol in beer production, some could produce ∼4% ethanol (v/v) ^22–24^. Beers brewed with *T. delbrueckii* that were able to produce moderate amounts of ethanol also showed high production of fruity flavours, potentially desirable in brewing ^22,23^. *T. delbrueckii* possesses various qualities, including low ethanol production and interesting floral flavours, that would make it suitable for the production of low-alcohol beers ^21,22^. Other non-*Saccharomyces* yeasts have been tested for potential roles in producing low-alcohol beers, including *Saccharomycodes ludwigii, Zygosaccharomyces rouxii*, and *Candida shehatae* ^31,32^. Producing beer using a single strain of wild yeast is typically difficult, as previously characterised wild yeast are frequently unable to utilise maltose and maltotriose ^21,33,34^, leaving the fermented beer with large amounts of normally fermentable sugars and low levels of ethanol, and hence prone to contamination. This problem is frequently overcome by co-inoculation with *S. cerevisiae* at the start or midpoint of fermentation ^21,33,35–37^. This provides the opportunity for flavour production by the wild yeast but ensures the fermentation will proceed to completion.

Brewing with spontaneous fermentation styles can produce beer with different and interesting flavours, but at a cost of decreased predictability and reproducibility. Fermenting with individual strains of wild yeast can provide robustness and reproducibility, but generally results in lower ethanol production. Although co-pitching with *S. cerevisiae* can provide efficient fermentation it can also remove or hide the nuanced flavours provided by the wild yeast. Here, we isolated diverse non-conventional yeast for the production of unique and unusual beers, aiming to achieve the predictability and reproducibility gained by using single, tested strains of yeast. We use proteomics, genomics, and metabolomics to characterise a few candidate strains, and to guide optimal process design to allow beer production without co-pitching with conventional brewing yeast, *S. cerevisiae*.

## Methods

### Collection and Isolation of Wild Yeast

Plant material (leaves, flowers, and bark) were collected, small (5-10 cm) sections were placed in 20 mL of rich media (YPD + 8% ethanol) to select for ethanol tolerant yeast and inhibit the growth of bacteria. Cultures were incubated at room temperature (RT) for 5 days, the liquid culture vortexed to resuspend cells, and 200 μL of each was plated on to YPD agar and incubated at RT for 3 days. Yeast isolates that grew were screened by assessing colony and cell size and cell shape. Yeast-like isolates were re-streaked onto YPD agar at RT to ensure clonality, grown in liquid YPD at RT and stored in glycerol stocks at -80°C. Three *S. cerevisiae* strains were used as fermentation controls: BY4741, *MATa* S288C-derivative laboratory strain; SafAle US05 american ale yeast (Fermentis); and M20, Bavarian wheat yeast (Mangrove Jacks).

### Identification by ITS1 (Internal Transcribed Sequence) Region

Identification of yeast species was performed initially by PCR of the internal transcribed spacer one (ITS1) region as previously described for genus and species level taxonomy of fungi ^6,38–41^. The primers for ITS1 region amplification were 5’–TTTCCGTAGGTGAACCTGC–3’ (forward) and 5’– GCTGCGTTCTTCATCGATGC–3’ (reverse), as described ^42^, with minor alterations to the forward primer to better match the Tm of the reverse primer. For Sanger sequencing, both primers were shortened to lower the Tm in line with the annealing temperature: 5’– AGGTGAACCTGC–3’ and 5’–TTCTTCATCGATGC–3’.

### Growth Assays

Growth of individual yeast isolates were tested in specified media or Dry Malt Extract (DME) wort, with 1.040 gravity DME, and 16.4 International Bitterness Units (IBUs) by the addition of Magnum Hops (which have an alpha acid composition of 12 – 14%). DME wort was inoculated with yeast at a starting OD_600nm_ of 0.1, equivalent to ∼1.28 × 10^6^ cells/mL. Ferments were tracked by weight loss due to glucose consumption and CO_2_ and ethanol production, an indicator of fermentation extent ^43^. Weight loss due to evaporation was controlled for by subtracting the weight loss of uninoculated wort.

### Headspace GC-MS for Ethanol Production

Samples were diluted 1:50 with H_2_O with 0.05% isopropanol as an internal standard. Ethanol was quantified using external calibration to multi-point standard curves with R^2^ values > 0.98. Chromatography was performed with a Rxi-624Sil 3.0 µm 30 m x 0.53 mm column (Restek). Headspace GC-MS was performed as described ^44^, with a total run time of 20 min per sample.

### Sugar and Amino Acid Quantification by Multiple Reaction Monitoring (MRM) LC-MS/MS

Fermented samples were prepared and measured as previously described ^45^. Samples were filtered and diluted 1:1000 with H_2_O. Sugars (glucose, maltose, and maltotriose) and amino acids (L-Serine, L-Proline, L-Valine, L-Threonine, L-Leucine, L-Isoleucine, L-Aspartic Acid, L-Lysine, L-Glutamic Acid, L-Methionine, L-Histidine, L-Phenylalanine, L-Arginine, L-Tyrosine, and L-Cystine) were quantified using external calibration to multi-point standard curves with R^2^ values > 0.99. MRM Targets are previously described ^45^, adapted from previous reports ^46,47^.

### Pressure Sensitivity

Pressure sensitivity of selected isolates was tested by inoculating 5 mL YPD with a starting OD_600nm_ of 0.1 in capped 10 cc/mL syringes with minimal headspace. These syringes were then incubated upright at RT with or without weight applied to the plunger, creating the equivalent of 2 or 1 atm (atmospheric pressure) environments, respectively.

### Whole Cell Sample Preparation

Proteins from yeast whole cell extracts were prepared for LC-MS/MS proteomics as previously described ^48^. Briefly, harvested cells were resuspended in non-denaturing buffer with protease inhibitors, lysed by bead beating, cysteines were reduced/alkylated, and centrifuged. Proteins from the soluble fraction were then desalted by precipitation with methanol/acetone. Finally, proteins were resuspended in 50 mM ammonium bicarbonate and digested with trypsin.

### Mass Spectrometry

Peptides were desalted with C18 ZipTips (Millipore) and measured by LC-ESI-MS/MS using a Prominence nanoLC system (Shimadzu) and TripleTof 5600 instrument with a Nanospray III interface (SCIEX) as previously described (36). Approximately 1 µg or 0.2 µg desalted peptides, as estimated by ZipTip binding capacity, were injected for data dependent acquisition (DDA) or data independent acquisition (DIA), respectively. Data Dependent Acquisition and Data Independent Acquisition LC-ESI-MS/MS were performed as previously described ^49^, with a total run time of 70 min per sample.

### Mass Spectrometry Data Analysis

Peptides and proteins were identified using ProteinPilot 5.0.1 (SCIEX), searching against either S288C *S. cerevisiae* (yeast) (Saccharomyces Genome Database (SGD), downloaded December 2017; 6,726 proteins) or proteins from whole genome sequences, as described below, with settings: sample type, identification; cysteine alkylation, acrylamide; instrument, TripleTof 5600; species, none; ID focus, biological modifications; enzyme, trypsin; search effort, thorough ID. The proteins identified by ProteinPilot were used to generate ion libraries for SWATH analysis. The abundance of peptides and proteins were determined using PeakView 2.1 (SCIEX) as previously described ^50,51^. PeakView 2.1 (SCIEX) was used for analysis with the following settings: shared peptides, allowed; peptide confidence threshold, 99%; false discovery rate, 1%; XIC extraction window, 6 min; XIC width, 75 ppm.

Protein-centric analyses was performed as previously described ^52^, protein abundances were re-calculated using a strict 1% FDR cut-off ^53^. Normalisation was performed to either the total protein abundance in each sample or to the abundance of trypsin self-digest peptides, as previously described ^51^. Principal component analysis (PCA) was performed using Clustvis 0.0.0.9 ^54^ in R. Protein and sample clustering was performed using Cluster 3.0 ^55^, implementing a hierarchical, uncentered correlation, and complete linkage. For statistical analysis, PeakView output was reformatted as previously described ^53^ and significant differences in protein abundance were determined using MSstats (2.4) ^56^ in R, with a significance threshold of p = 10^-5^ ^57^. Gene ontology (GO) term enrichment was performed using GOstats (2.39.1) ^58^ in R, with a significance threshold of p = 0.05 ^57^. Large GO term lists were reduced by summarising and removing redundances using REVIGO ^59^ in R.

### Whole Genome Sequencing

Whole genome sequencing of wild yeast isolates was performed via a shotgun metagenomic pipeline using an Illumina NextSeq550 with 100 x coverage. Raw reads were processed with Trimmomatic, assembled with Shovill, and quality checked using Benchmarking Universal Single-Copy Orthologs (BUSCO) using the Saccharomycetes lineage dataset. Assemblies were also evaluated with Quast (ver. 5.02; --fungus), using the reference genome sequences and features for the *S. cerevisiae* model strain S288C (https://www.yeastgenome.org/strain/s288c) and the *T. delbrueckii* type strain (https://www.ncbi.nlm.nih.gov/assembly/GCF_000243375.1/). RepeatMasker was used to identify and mask repeat regions in the assemblies. Protein coding genes within the masked assemblies were predicted with GeneMark-ES. Protein sequences were functionally annotated by BLAST with DIAMOND blastp against the SwissProt database.

### Data availability

The mass spectrometry proteomics data have been deposited to the ProteomeXchange Consortium via the PRIDE partner repository ^60^.

## Results

### Capturing and isolating unique yeast from diverse flora

We set out to isolate, identify, and assess the fermentation profile of wild yeast for brewing beer. Here, we outline a collection, isolation, storage, and fermentation workflow for wild yeast characterisation (Fig. 1A). We collected samples (flowers, leaves, or bark) from 47 plants at The University of Queensland’s St Lucia campus in Brisbane, Australia (Fig. 1B and C). We cultured plant samples in YPD + 8% ethanol for 5 days, resuspended yeast cells, plated a fraction to YPD agar, and incubated until colonies grew (Fig. 1A and 1D). Yeast-like colonies were re-streaked onto YPD to guarantee single clonal colonies were selected, and morphology and microscopy were assessed to confirm isolates were non-filamentous yeast (Fig. 1A). 11 of the plant material samples yielded yeast, with a total of 30 unique isolates.

**Figure 1.**
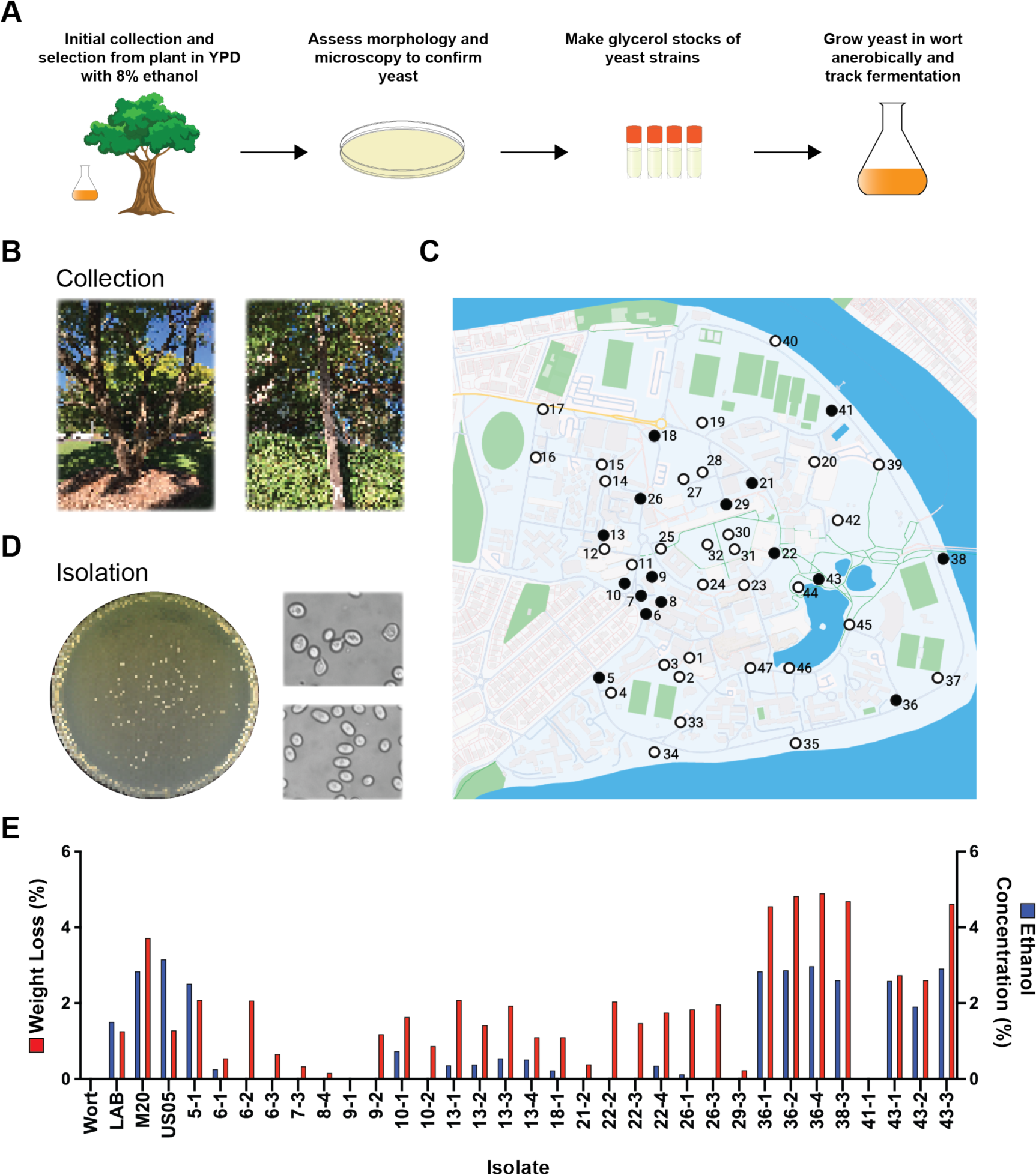
Collection, isolation, and fermentation of wild yeast isolates. **(A)** Overview of the workflow for collection, isolation, and initial characterisation of wild yeast isolates. **(B)** Example collection sites. **(C)** Map showing location of sampling and highlighting locations of successful yeast isolation. White, no yeast isolated; black, yeast isolated. **(D)** Example of isolate plate morphology and microscopy. **(E)** Concentration of ethanol produced (blue) and total weight loss (red) of yeast strains. Control ferments labelled; laboratory yeast - BY4741, brewing yeast – US05, and brewing yeast – M20, other lines represent individual wild yeast ferments.

Once individual yeast strains had been isolated, we assessed their growth in wort to determine their suitability for brewing beer (Fig. 1E). To measure fermentation extent, we inoculated wort with equal amounts of yeast, and tracked the weight loss of fermentations across 25 days (Fig. 1E). Ferments were tracked by weight loss due to glucose consumption and CO_2_ and ethanol production. The control *S. cerevisiae* ferments (BY4741, M20, and US05) had a total weight loss of 1.26%, 3.71%, and 1.28%, respectively and produced 1.50%, 2.84%, and 3.16% ethanol (v/v), respectively. Six wild yeast isolates had a lower total weight loss than US05 and BY4741 and 15 isolates produced less ethanol than BY4741 (1.50%) (Fig. 1E). Three isolates had a higher total weight loss and produced more ethanol than BY4741 but less than US05 and M20 (Fig. 1E). Interestingly, five wild yeast isolates (36-1, 36-2, 36-4, 38-3, and 43-3) had a higher total weight loss than all three control ferments and produced between 2.58 – 2.91% ethanol (v/v), similar to M20 which produced 2.84% ethanol (Fig. 1E). When comparing weight loss with ethanol production, some wild yeast produced little ethanol while losing substantial amounts of weight, suggesting that such isolates are likely either producing other fermentation end products and/or do not produce ethanol as efficiently as *S. cerevisiae* strains.

We successfully identified 21 of 30 wild yeast isolates by Sanger sequencing of the ITS1 region (Supplementary Table 1), where we defined the identity of each isolate by the species that best matched its ITS1 sequence including primer regions by BLAST search. We examined the relationship between our 21 identifiable isolates and selected published yeast species by multiple sequence alignment and phylogenetic analysis of ITS1 regions using Clustal Omega (Fig. 2). We found that six isolates were *T. delbrueckii* and another two were able to be placed within the *Torulaspora* genus but could not be identified to species level (Supplementary Table 1). All the *Torulaspora* strains we identified clustered within the *Torulaspora* clade, with the unclassified *Torulaspora* strains (36-1, 36-2, 36-4, and 38-3) clustering in a sub-clade with *Torulaspora* strains isolated from soil (AY046186, KF300900, AY046188, and KF300901) (Fig. 2 and Table S2). 13-1, 13-2, 13-3, and 13-4 sat with a *T. delbrueckii* isolated from an unspecified source, between the soil sub-clade and a sub-clade consisting of clinical and fermented product isolates (Fig. 2 and Table S2).

**Figure 2.**
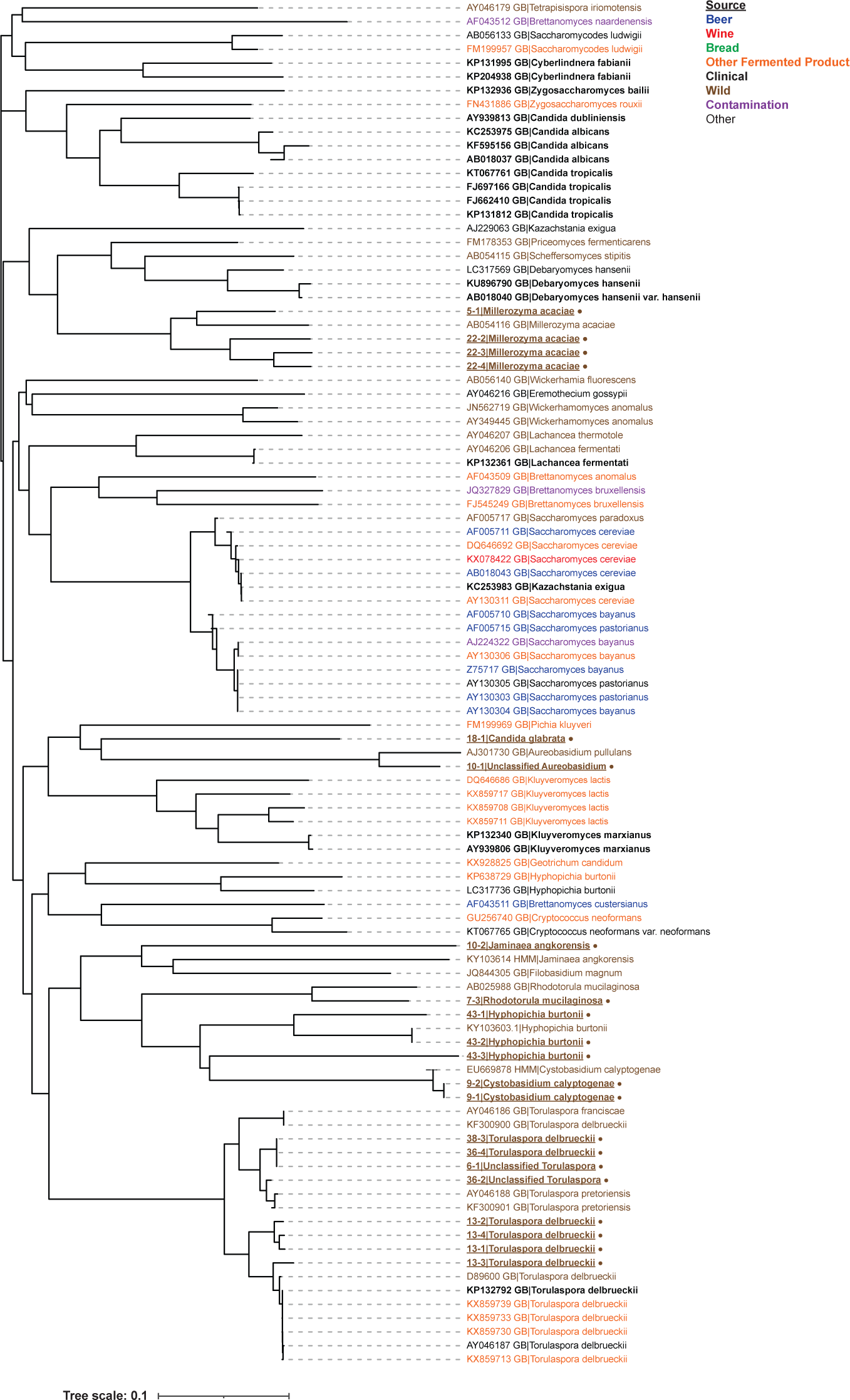
Phylogenetic tree of ITS1 region of all identified yeast isolates and select published strains. Phylogenetic tree calculated using Clustal Omega of ITS1 region of isolated wild yeast and select strains and species within the order *Saccharomycetales*. Strains isolated in this work are underlined, bold, and asterisked. For strain classification see Supplementary Table S2.

We identified four isolates (5-1, 22-2, 22-3, and 22-4) as *Millerozyma acacia*, all of which clustered in a clade with AB054116, a *M. acacia* isolated from a larva of a jewel beetle (Fig. 2 and Table S1 and S2). We identified three isolates (43-1, 43-2, and 43-3) as *Hyphopichia burtonii* which clustered with a wild isolate of *H. burtonii* (KY103603.1) (Fig. 2 and Table S1 and S2). We identified two isolates (9-1 and 9-2) as *Cystobasidium calyptogenae* isolates, one isolate (7-3) as *Rhodotorula mucilaginosa,* one isolate (10-2) as *Jaminaea angkorensis*, and one isolate (10-1) as an unclassified species of *Aureobasidium*, all of which clustered with related species, most of which had been isolated from the wild (Fig. 2 and Table S1). We also identified one isolate (18-1) as *Candida glabrata*, which interestingly did not cluster with *Candida* species but rather with *Pichia* and *Aureobasidium* (Fig. 2 and Table S1).

### Utilisation of sugars and amino acids by select top-performing strains

Our initial fermentation assay showed that a select few wild yeast isolates were able to efficiently produce ethanol at a similar level to brewing yeast (Fig. 1). To better understand any stresses or limitations the wild yeast strains might face in metabolising nutrients in a brewing environment, we investigated the sugar and amino acid profiles post-fermentation in wort. We compared: two high-performing unclassified *Torulaspora* isolates, 36-1 and 38-3; brewing yeast US05; and 13-2, a poor fermenting *T. delbrueckii* isolate. MRM analysis showed that the fermentable sugar profiles post-fermentation were similar between 36-1, 38-3, and US05 (Fig. 3A). Starting wort contained 9.26 mg/mL glucose and after fermentation there was 1 mg/mL present when fermented with US05 or 13-2, and ∼ 0 mg/mL when fermented with 36-1 or 38-3 (Fig. 3A). Starting wort contained 57.83 mg/mL maltose, and after fermentation there was 28.64 mg/mL, 63.56 mg/mL, 39.90 mg/mL, or 25.48 mg/mL maltose when fermented with US05, 13-2, 36-1, or 38-3, respectively (Fig. 3A). There was ∼15 mg/mL maltotriose present in both pre-fermentation wort and after fermentation regardless of yeast strain. Together, this is consistent with all yeasts efficiently using glucose; US05, 36-1, and 38-3, but not 12-3, partially using maltose; and none of the yeasts efficiently using the larger maltotriose (Fig. 3A). We then grew these strains on rich media with glucose or maltose as the sole carbon source and found that all strains except 13-2 were able to grow with maltose as the sole carbon source (Fig. 3B). These results showed that 36-1 and 38-3 were able to grow on maltose and glucose just as well as US05, while 13-2 was unable to utilise maltose, and none of the strains were able to utilise maltotriose (Fig. 3A and B). Yeasts need to be able to grow with glucose or maltose as sole carbon sources to be able to efficient ferment wort, and the inability of most wild yeasts to metabolise maltose potentially limits fermentation and would lead to either a low alcohol beer or the need to co-inoculate with brewing *S. cerevisiae* strains ^21,33,35–37^. It was therefore interesting and striking that both 36-1 and 38-3 could use maltose as a sole carbon source and grow efficiently in wort, a rare trait amongst *Torulaspora* ^21,22^.

**Figure 3.**
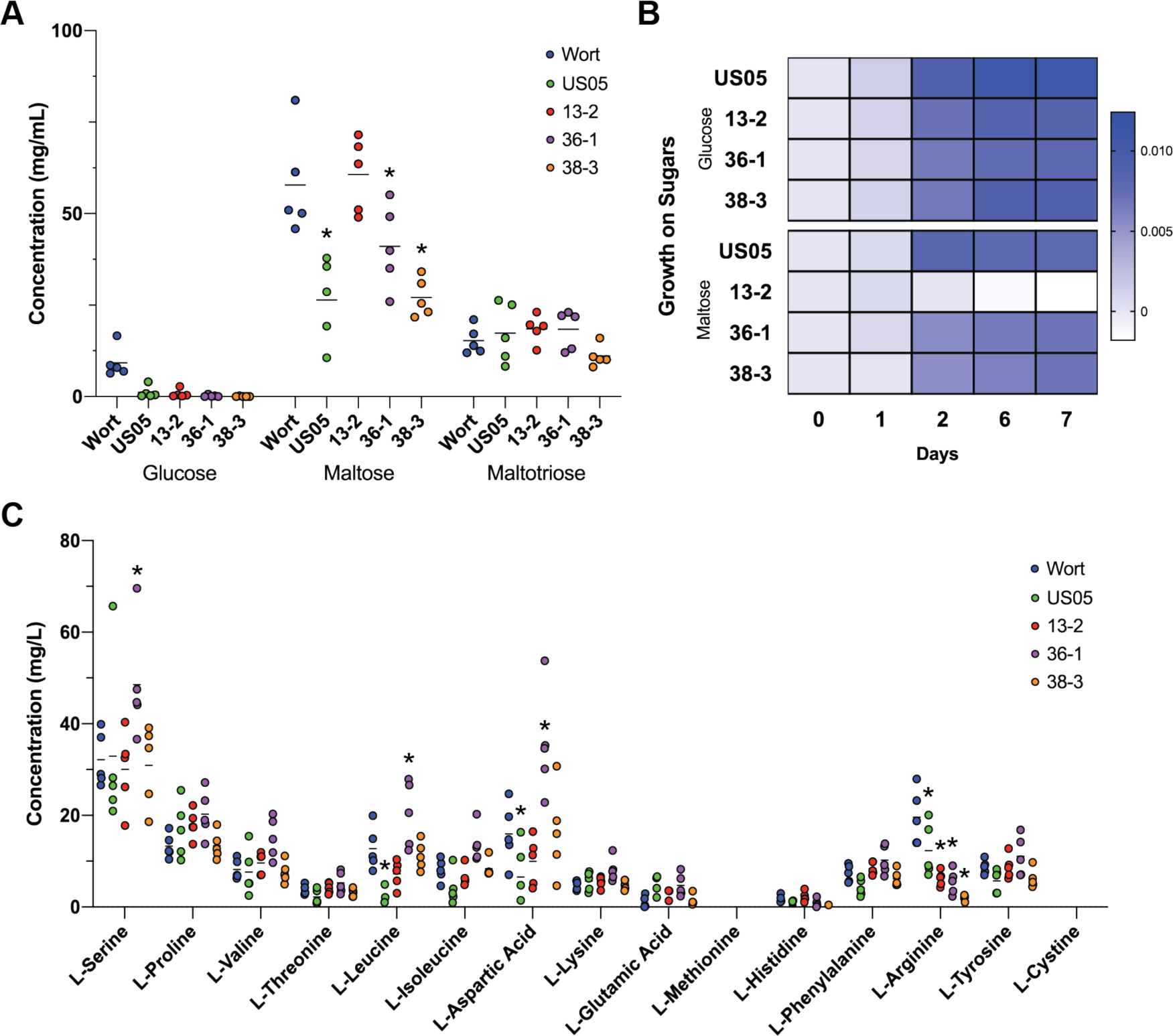
Utilisation of sugars and amino acids by candidate wild yeast isolates. **(A)** Glucose, maltose, and maltotriose quantified in uninoculated wort and after fermentation by wild yeast isolates or US05 brewing yeast. *, p < 0.05 compared with wort. **(B)** Utilisation of glucose and maltose as sole carbon sources, shown as proportion of weight loss over time compared to original weight. **(C)** Amino acids quantified in uninoculated wort and after fermentation by wild yeast isolates or US05 brewing yeast. *, p < 0.05 compared with wort.

Like all organisms, yeast require a source of nitrogen for growth. Generally, the preferred source of nitrogen for yeasts is from amino acids present in the growth medium. We therefore investigated how the wild yeast strains utilised the amino acids present in wort. We were able to measure 13 free amino acids in wort using MRM LC-MS/MS and found that all three wild yeast strains as well as US05 showed similar amino acid utilisation preferences (Fig. 3C). As all of the *Torulaspora* strains were able to utilise amino acids with similar efficiency to US05 (Fig. 3C), we concluded that amino acid availability or preference was unlikely to present difficulties in beer fermentation using these isolates.

### Failed fermentations at larger scales highlight possible growth limiting factors

As small-scale fermentation showed no apparent limitations with the use of 36-1 or 38-3 for beer production from wort, we proceeded with upscaling fermentation towards industrial scale. We performed fermentation of wort with 36-1 at a range of scales (5 L, 100 L, and 1000 L) all with a starting specific gravity of ∼1.040 (Fig. 4A). Specific gravity (SG) relates to the relative density of a solution compared to water, and roughly indicates the amount of fermentable sugar present. Measuring specific gravity at the beginning (original gravity, OG) and end of fermentation (final gravity, FG) can be used to determine the amount of alcohol present and fermentable sugar remaining. Typically, a finished beer has an FG of 1.010 or below. The ferments at both 5 L and 100 L scales both finished with an FG of ∼ 1.018 (2.89 % ABV), appropriate for a mid-strength beer (Fig. 4A). We then performed two ferments at 1000 L which surprisingly and unfortunately finished with FGs of 1.030 and 1.037, indicating incomplete fermentation (Fig. 4A). These results suggested that fermentation with 36-1 at 1000 L was not able to achieve the same efficient growth as at a smaller scale. If beer was packaged with this incomplete fermentation, the high levels of residual fermentable sugars would create a substantial risk of infection or re-fermentation.

**Figure 4.**
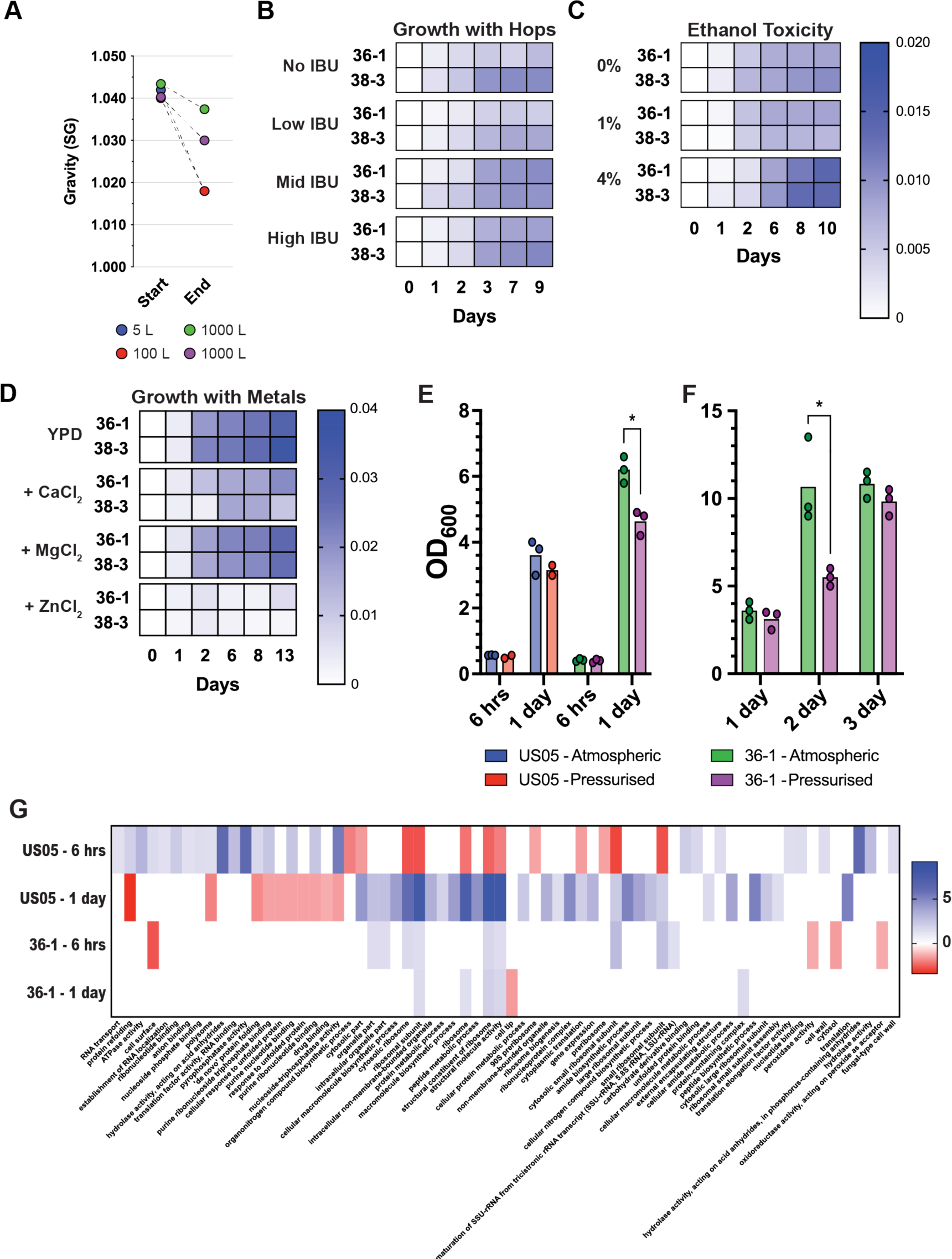
Discovering fermentation parameters that limit growth at industrial scale. **(A)** Original and final gravity (OG and FG, respectively) of different scale ferments. All ferments used strain 36-1. **(B)** Hop toxicity of candidate isolates in wort with different IBUs, measured as proportion of weight loss over time compared to original weight. **(C)** Ethanol toxicity of candidate isolates in YPD with different concentrations of ethanol, measured as proportion of weight loss over time compared to original weight. **(D)** Growth in YPD with additional 10 mM CaCl_2_, MgCl_2_, or ZnCl_2_ measured as proportion of weight loss over time compared to original weight. **(E)** Growth of 36-1 and US05 at atmospheric pressure (1 atm) and pressurised (2 atm). OD_600_ measured 6 h and 1 day post-inoculation. Values, mean. *, p < 0.05. **(F)** Growth of 36-1 at atmospheric pressure (1 atm) and pressurised (2 atm) across 3 days, with intermittent release of pressure. Values, mean. *, p < 0.05. **(G)** Significantly enriched GO terms in proteins significantly more abundant (blue) or less abundant (red) in cells grown while pressurised compared to atmospheric pressure from (E). Values shown as −log_2_ of Bonferroni corrected p-value for GO terms which were significantly enriched (p < 0.05).

To better understand the factors that limited growth at a commercial scale we tested the growth of candidate wild yeasts 36-1 and 38-3 in the presence of hops, ethanol and different metals (CaCl_2_, MgCl_2_, or ZnCl_2_), components of wort which can inhibit growth of some bacteria and yeast. We first assessed the effect of increasing amounts of hops/bitterness (international bitterness units, IBU) on the growth of 36-1 and 38-3 (Fig. 4B). Both 36-1 and 38-3 grew better in mid and high IBU wort compared to low or no IBU wort, having a greater total weight loss (Fig. 4B). This showed that the presence of hops was not a growth limiting factor in fermentation for 36-1 or 38-3. The improved fermentation in wort with more hops is potentially linked to the small amounts of amyloglucosidase, α-amylase, β-amylase, and limit dextrinases in hops, which together can increase the concentration of fermentable sugars ^61^. Next, we assessed the toxicity of exogenous ethanol to 36-1 and 38-3, which showed that neither strain’s growth was impeded by the presence of ethanol at concentrations relevant to beer fermentation (Fig. 4C). Finally, we assessed the effect of different metals on 36-1 and 38-3 growth. Addition of ZnCl_2_ dramatically impeded the growth of both 36-1 and 38-3, while CaCl_2_ slightly reduced the growth of both isolates (Fig. 4D). Addition of MgCl_2_ had no effect on the growth of 36-1 and 38-3 (Fig. 4D). Together, this demonstrated that ethanol and hops at concentrations relevant to those found in wort and beer did not limit growth of 36-1 or 38-3. While ZnCl_2_ did impede fermentation, zinc is rarely found at high concentrations in wort.

One of the most pronounced differences between small- and large-scale fermentation is the hydrostatic pressure created by the physical scale of fermenter vessels. To better understand the effect of fermentation scale and hydrostatic pressure on yeast physiology and growth, we grew US05 and 36-1 at atmospheric pressure (1 atm) and under one additional atmosphere of pressure (2 atm), equivalent to the pressure at the bottom of the liquid in a 10 m tall fermenter. We found that additional hydrostatic pressure did not influence growth of US05, but that the growth of 36-1 was significantly hindered by the additional hydrostatic pressure after 1 day of fermentation (Fig. 4E). This is consistent with the adaptation of the commonly used brewing yeast US05 to the brewing environment. As 36-1 and US05 behaved differently when grown under additional pressure, we investigated the impact of this additional pressure on the whole cell proteomes of US05 and 36-1 after 6 h and 1 day of growth (Fig. 4E and G). We performed MSstats comparison of the proteomes of yeast (both US05 and 36-1) grown at atmospheric pressure and under additional pressure, followed by GO term enrichment analysis on proteins significantly higher or lower in abundance at atmospheric pressure or under additional pressure (Fig. 4G, Table S3, and Table S4). We found that US05 had a significant change in GO term enrichment from 6 h to 1 day (Fig. 4G). Most interestingly, US05 under pressure at 6 h showed an enrichment of proteins involved with cell wall processes, likely related to the stresses of additional pressure (Fig. 4G). However, this enrichment of cell wall processes was no longer seen in US05 after 1 day (Fig. 4G). Next, we investigated the proteomic response of 36-1 to growth under additional pressure and found that 36-1 had few GO terms enriched between atmospheric pressure and under additional pressure at 6 h, and after 1 day had even fewer GO terms enriched (Fig. 4G). The enrichment of cell wall related GO terms at 6 h of US05 followed by the loss of enrichment at 1 day hinted at the mechanisms that underlay the ability of US05 to adapt to growth in the presence of additional pressure. The absence of such a response in 36-1 may partially explain its failure to grow at large scale. We next tested if growth was temporarily or permanently stalled by this additional pressure. We grew 36-1 under additional pressure for 3 days (Fig. 4F). At 1 day we found no difference in growth between atmospheric pressure and additional pressure in either 36-1 (Fig. 4F). At 2 days we found that the growth of 36-1 was significantly hindered by additional pressure (Fig. 4F). Finally, we found that at 3 days, the growth of 36-1 recovered to be consistent with or without additional pressure (Fig. 4F). The growth of yeast under similar pressures to those found in the brewing process is highly relevant when investigating wild yeast for brewing. These results also suggested that while additional pressure did slow the growth of wild yeast, the microbes could overcome this pressure and still effectively finish fermentation.

### Modification to fermentation conditions to alleviate stress and promote growth

As we did not identify a limiting factor(s) that substantially affected the growth of wild yeast isolates, we investigated if fermentation conditions could be manipulated to encourage yeast viability. To this end we tested the growth of 36-1 and 38-3 at higher temperature, with additional glucose, or with additional amino acids (Fig. S2). Fermentation performance was slightly improved with the addition of glucose and amino acids, while the increased temperature did not affect fermentation (Fig. S2). These results suggested that addition of extra glucose or amino acids could benefit fermentation performance and could be considered to improve commercial fermentation, but that they were unlikely to substantially improve fermentation performance.

### Dual stresses limit wild yeast growth at industrial scale

As independent application of potential stresses associated with growth at industrial scale did not substantially affect the growth of wild yeast isolate 36-1 (Fig. 3), we next investigated if it was the combination of multiple stresses which caused slow growth. The two key stresses associated with beer production at industrial scale are the presence of maltose as the key carbon source and the high hydrostatic pressure. While both of these stresses slowed the growth of 36-1 when applied independently, the yeast could still complete fermentation (Fig. 3 and 4). We therefore explored how the combination of these stresses impacted the growth of 36-1. We grew yeast on media with either glucose or maltose as the sole carbon source, and at atmospheric pressure or with an additional atmosphere of pressure applied. Consistent with our previous results, we found that independent application of each stress did not affect growth (Fig. 5A). However, simultaneous application of both maltose and pressure stress significantly hindered growth compared to cultures grown with neither stress or only one of the two stresses (Fig. 5A). As it was only the combination of maltose and pressure stresses that limited growth, we tested if relieving one of the stresses would rescue growth. Glucoamylase digests maltose to glucose, and we found that addition of glucoamylase to media with maltose as the carbon source indeed rescued the growth of 36-1 yeast even in the presence of pressure stress. (Fig. 5A).

**Figure 5.**
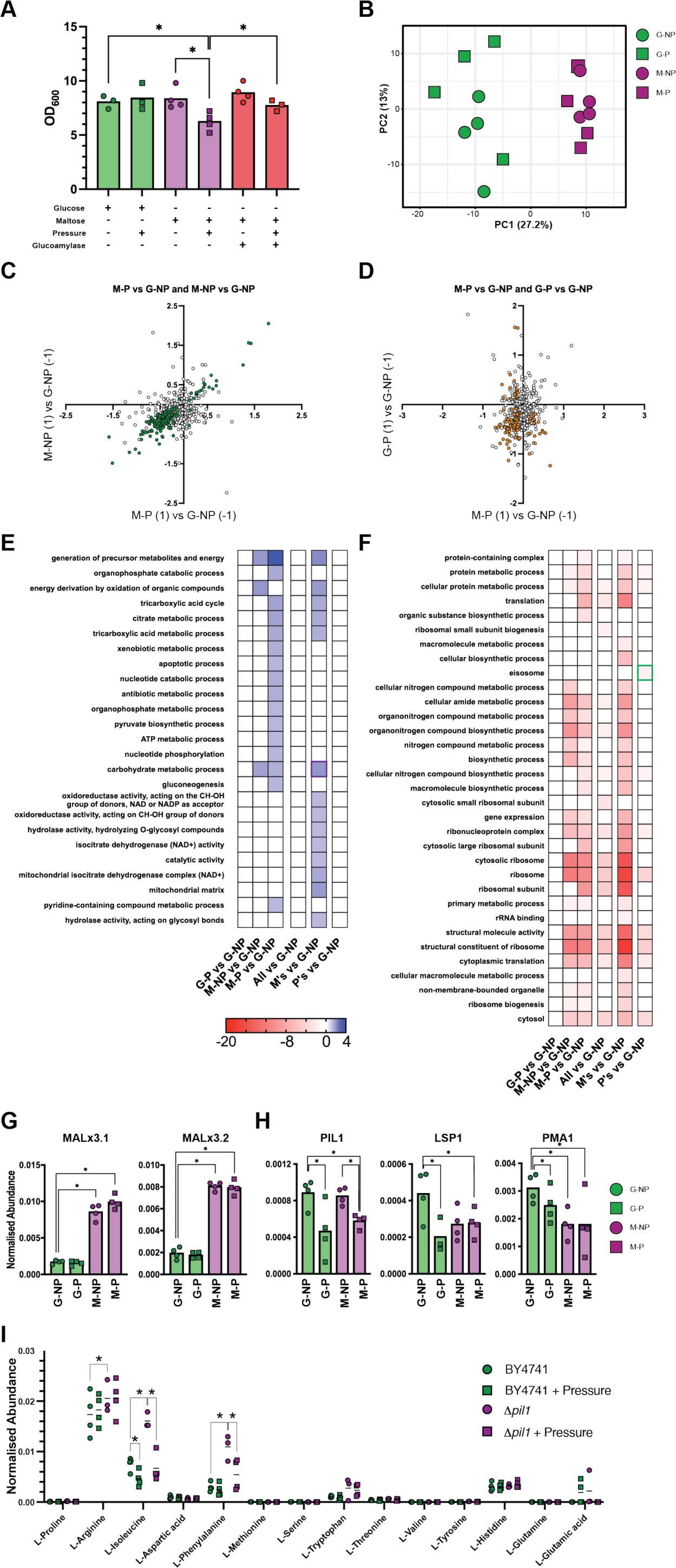
Dual stresses limit wild yeast growth. **(A)** Growth of 36-1 in YPD (Glucose) and YPM (Maltose) at atmospheric pressure (1 atm) and pressurised (2 atm), and with addition of glucoamylase. OD_600_ measured 72 h post-inoculation. **(B)** PCA of whole cell proteome of 36-1 under different growth conditions: G-NP (glucose – atmospheric pressure), G-P (glucose – pressurised), M-NP (maltose – atmospheric pressure), and M-P (maltose – pressurised). The first component (x-axis) accounted for 27.2% of the total variance and the second (y-axis) 13%. **(C)** Scatterplot of the Log2FC between M-P and G-NP (x axis) and Log2FC between M-NP and G-NP (y-axis). Clear circles, not significantly different in both comparisons; orange circles, significantly different in both comparisons (p < 10^-5^) **(D)** Scatterplot of the Log2FC between M-P and G-NP (x axis) and Log2FC between G-P and G-NP (y-axis). Clear circles, not significantly different in both comparisons; green circles, significantly different in both comparisons (p < 10^-5^). **(E)** GO terms enriched in proteins significantly more abundant in each condition labelled compared to G-NP. Purple box highlights GO term of interest and associated proteins in (**G**). **(F)** GO terms enriched in proteins significantly less abundant in each condition labelled compared to G-NP. Values shown as −log2 of Bonferroni corrected p-value for GO terms which were significantly enriched (p < 0.05). Green box highlights GO term of interest and associated proteins in **(H)**. GO Terms shown are non-redundant and summarised calculated by REVIGO. **(G)** Normalised abundance of sphingolipid long chain base-responsive protein Pil1, sphingolipid long chain base-responsive protein Lsp1, and Plasma membrane ATPase 1 Pma1. Bar, mean. *, p < 10^-5^. **(H)** Normalised abundance of two alpha-glucosidases (MALx3.1 and MALx3.2). **(I)** Normalised abundance of measured amino acids. *, p < 0.05.

To gain insights into the mechanistic consequences of and responses to dual maltose and pressure stress in 36-1, we investigated the impact of maltose and pressure stress on the yeast whole cell proteome (Table S5 and S6). We grew 36-1 yeast in four conditions: glucose-atmospheric pressure (G-NP), glucose-pressurised (G-P), maltose-atmospheric pressure (M-NP), and maltose-pressurised (M-P), and performed proteomic analysis of whole cell extracts. PCA revealed that growth in maltose substantially changed the proteome of 36-1, while additional pressure had little effect (Fig. 5B). To identify the proteomic changes in 36-1 grown in maltose as the carbon source that were consistent at both pressures, we plotted the Log2FC of protein abundance in M-NP and M-P, both compared to G-NP, with only significantly different proteins coloured (Fig. 5C). We found a linear relationship, highlighting that the changes in the proteome were primarily due to maltose stress, with several proteins more abundant in both maltose conditions and many proteins significantly less abundant in both maltose conditions, compared to G-NP. We next performed GO term enrichment on proteins significantly more abundant in the maltose conditions compared with G-NP, which identified a noticeable enrichment of carbohydrate metabolic process (Fig. 5E). This was associated with two maltase activator proteins, the transcription factor for the maltose transporter and maltase (Fig. 5G). The enrichment and significant increase of maltose related proteins is consistent with 36-1 changing its sugar utilisation pathways to maltose when grown in media containing only maltose as a carbon source.

Next, we identified the proteomic changes in 36-1 grown under additional pressure that were shared across both carbon sources. We plotted the Log2FC of protein abundance in M-P and G-P, both compared to G-NP, with only proteins with significantly different abundance coloured (Fig. 5D). This comparison did not show a strong linear relationship, again consistent with changes in carbon source dominating the proteomic response of 36-1, rather than changes in pressure. We also found that no proteins were significantly more abundant in both pressure conditions compared to glucose without pressure. However, we did identify a suite of proteins that were significantly less abundant in both pressure conditions compared to glucose without additional pressure. GO term enrichment of these proteins that were significantly less abundant in both pressure conditions compared with G-NP showed a noticeable enrichment of GO terms related to metabolic processes and ribosomes (Fig. 5E and F). The abundance of these classes of proteins are typically correlated with growth rate, consistent with their lower abundance in cells under pressure stress ^62,63^. Interestingly, we also identified an enrichment of proteins associated with eisosomes in cells grown under pressure stress (Fig. 5F). Eisosomes are membrane invaginations at the cell surface that harbor Amino Acid-Polyamine-Organocation (APC) transporters and help maintain plasma membrane integrity ^64–66^. When they are not needed for active amino acid uptake, APCs are maintained within eisosomes, where they are inactive, but from where they can be readily released into the neighbouring plasma membrane when needed ^64,67,68^. In our proteomic data, three proteins with lower abundance in cells under pressure stress were specifically associated with eisosomes: sphingolipid long chain base-responsive protein Pil1, sphingolipid long chain base-responsive protein Lsp1, and Plasma membrane ATPase 1 (Pma1) (Fig. 5H). Pil1 and Lsp1 are membrane-associated proteins that physically help form eisosomes ^69^, while Pma1 is a proton pump that drives the active transport of nutrients, including through APC transporters ^64,66^.

The proteomic changes we observed in 36-1 suggested that pressure stress relevant to fermentation at industrial scale was mechanistically associated with defects in amino acid uptake. Specifically, we observed that pressure was associated with low abundance of the homologous proteins Pil1 and Lsp1, key proteins required for the formation of eiososomes. To test if loss of Pil1/Lsp1 caused changes in amino acid uptake, we compared the intracellular amino acid profile in the genetically tractable wild type *S. cerevisiae* yeast and Δ*pil1* yeast, with and without additional hydrostatic pressure. Growth with additional pressure caused a significant decrease in the abundance of isoleucine in wild type yeast (Fig. 5I). However, loss of Pil1 caused a significant increase in the intracellular abundance of several amino acids, which was reversed in growth with additional pressure. The increased intracellular amino acid abundance in Δ*pil1* cells is consistent with the normal role of Pil1 in sequestering APCs in inactive forms in eisosomes, while the decreased amino acid abundance with additional pressure is consistent with high pressure inhibiting the amino acid transport activity of APCs ^64,66–68^. Together, these results support a model of response to pressure and maltose stress in 36-1. In this model, high pressure reduces the efficiency of amino acid uptake into cells, causing a regulated reduction in Pil1 abundance in order to increase the levels of active APCs at the plasma membrane. This homeostatic mechanism can rescue nutrient uptake when cells are grown on glucose. However, growth on the suboptimal carbon source maltose causes, amongst other changes, a reduction in the abundance of the glucose-regulated proton exporter Pma1. Critically, APCs require a Pma1-dependent proton gradient to import amino acids ^64^. Growth can therefore be maintained in the presence of either stress independently, but their combination causes a synergistic metabolic defect that inhibits growth. Some yeasts, such as commercial brewing yeasts, are able to overcome these combined stresses, either due to the inherent cell biology of *Saccharomyces* yeasts or to evolved features that have been acquired during domestication. However, the wild *Torulaspora* 36-1 yeast succumbs to the dual pressures inherent in fermentation at industrial scale.

### Successful industrial scale brewing

Given that the combination of maltose and pressure stresses hindered growth of 36-1, but that the addition of glucoamylase relieved maltose stress and allowed fermentation to continue to completion at laboratory scale, we next tested if the addition of glucoamylase was able to relieve stress and allow fermentation to complete at industrial scale. We therefore performed a 600 L fermentation of wort treated with glucoamylase (Figure 6).

**Figure 6.**
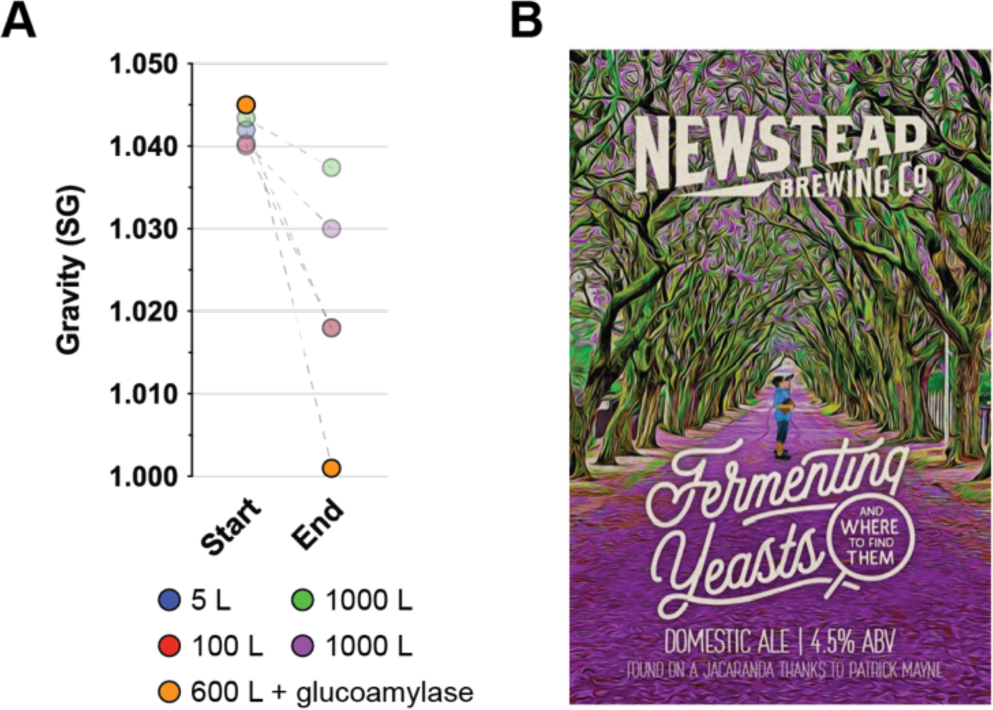
Commercial beer successfully brewed with 36-1. **(A)** Start and end gravity (SG) of different scale ferments. All ferments used strain 36-1. Orange indicates fermentation where glucoamylase was added **(B**) Decal of commercial beer made with Newstead Brewing Co.

The 600 L of wort had an OG of 1.045, and we inoculated it with 36-1 at a starting OD_600nm_ of 0.1 (Figure. 6A) together with commercial glucoamylase,. The ferment progressed efficiently and reached a FG of 1.001 (Figure. 6A). We then matured and packaged the beer (Figure 6B). This provided successful demonstration of the utility of the process interventions that we identified at laboratory scale when applied at industrial scale.

## Conclusions and Future Directions

We have developed and demonstrated a simple workflow for the isolation, identification, and fermentation assessment of wild yeast. Using this workflow, we were able to select high performing isolates, but found that they failed to efficiently ferment at industrial scale due to the combination of maltose and pressure stress. We used systems biology to uncover the mechanism underlying the synergistic metabolic defects caused by these dual stresses, and demonstrated that glucoamylase addition could relieve maltose stress to allow industrial scale fermentation using wild yeast. We see exciting potential in using our now established workflow on additional isolates to create a collection of strains of fermentation-capable wild yeast for beer brewing, to identify process or metabolic interventions to allow their use at industrial scale, and to better understand the fermentative, genomic, and metabolic diversity of wild yeast.

